# Coagulation Factor IIIa (*f3a*) Knockdown in Zebrafish Leads to Defective Angiogenesis and Mild Bleeding Phenotype

**DOI:** 10.1101/2021.12.21.473736

**Authors:** Saravanan Subramaniam, Jiandong Liu, Craig Fletcher, Ramani Ramchandran, Hartmut Weiler

**Affiliations:** Pulmonary Center, Department of Medicine, Boston University School of Medicine, Boston, MA, USA; Blood Research Institute, Blood Center of Wisconsin: part of Versiti, Milwaukee, WI, USA; UNC McAllister Heart Institute, University of North Carolina at Chapel Hill, NC, USA; Department of Pathology and Laboratory Medicine, University of North Carolina at Chapel Hill, NC, USA; Department of Pediatrics, Division of Neonatology, Medical College of Wisconsin, Milwaukee, WI 53226, USA

**Author notes:** **Corresponding author:** Saravanan Subramaniam, Ph.D., Department of Medicine, Boston University School of Medicine, Boston, MA, 02118, USA.

**Keywords:** Tissue factor, bleeding, angiogenesis, *f3a*

## Abstract

**Aim:** Tissue factor (TF), an initiator of the extrinsic coagulation pathway, is crucial for embryogenesis, as mice lacking TF are embryonically lethal (E10.5). This lethality may be attributed to defects in vascular development and circulatory failure, suggesting additional roles for TF in embryonic development beyond coagulation. In this study, we characterized the role of one of the TF paralogs (*f3a*) using a zebrafish model.

**Methods:** To understand the TF evolution across different species, we performed molecular phylogenetic and sequence homology analysis. The expression of *f3a* during embryonic developmental stages was determined by RT-PCR. Endothelial-specific transgenic lines of zebrafish (*flk1:egfp-NLS/kdrl:mCherry-CAAX*) was used to image the vascular development. The role of *f3a* during embryonic development was investigated by mRNA knockdown using Morpholinos (MO), an antisense-based oligonucleotide strategy. The *f3a* morphants were examined at 52 hpf for defects in morphological appearance, bleeding, and vascular patterning.

**Results:** Spatiotemporal expression of *f3a* by qPCR revealed expression in all developmental stages, suggesting that *f3a t*ranscripts are both maternally and zygotically expressed. High expression of *f3a* from 28 hpf to 36 hpf confirmed the role of in the development of the yolk sac, circulation, and fins. *f3a* MO-injected embryos showed morphological abnormalities, including shorter body lengths and crooked tails. O-dianisidine staining showed *f3a* MO-injected embryos exhibited bleeding in the trunk (5.44%) and head (9.52%) regions. Using endothelial-specific transgenic lines of zebrafish (*flk1:egfp-NLS/kdrl:mCherry-CAAX*), imaging of caudal vein plexus, which forms immediately following the onset of circulation and sprouting, showed a 3-fold decrease in *f3a* morphants versus controls at 48 hpf, suggesting a potential role for *f3a* in flow-induced angiogenesis.

**Conclusion:** *f3a* is essential for angiogenesis, in addition to its involvement in hemostasis.

**Key Points:**

- *f3a* knockdown by antisense morpholino oligonucleotide (MO)-approach shows bleeding phenotype.
- *f3a* is essential for angiogenesis, in addition to its involvement in hemostasis.

## Introduction

Tissue factor (coagulation factor III), a cell surface transmembrane glycoprotein receptor for coagulation factor VII/FVIIa, is a well-recognized trigger for the extrinsic coagulation cascade. The human tissue factor (TF) gene was cloned in late 1980s [1–3] and is localized on chromosome 1, p21-p22. TF is involved in inflammation, thrombosis, atherosclerosis, sepsis, tumor progression, embryogenesis [4, 5], and maintenance of vascular integrity in the placenta [6]. Disruption of the TF gene in mice is associated with embryonic lethality by E10.5 [7–9], consistent with the suggestion that humans cannot survive without TF [10]. Because TF-deficient mice are embryonically lethal, researchers who investigated embryonic developmental stages up to E10.5 have hypothesized that the lethality might be due to a defect in vascular development [7] and circulatory failure [8, 9]. Therefore, although TF is essential for embryonic development, its specific functions are not well defined due to embryonic lethality.

Angiogenesis is the growth of blood vessels from the existing vasculature and occurs throughout life, in health and disease conditions. The role of TF in tumor angiogenesis has been investigated for over two decades. One study concluded that the cytoplasmic domain of TF regulates the production of vascular endothelial growth factor (VEGF) [11–13]. Another study involving TF-deletion in mice concluded that yolk sac vessels of TF-deficient embryos were more fragile due to a deficit in mesenchymal cells/pericyte accumulation [7]. Embryonic lethality in mice has made investigations into the physiological role of TF in vascular development a bit challenging. Complementary to mouse model, the vertebrate zebrafish model system is used to investigate vascular development [14]. Zebrafish embryogenesis studies have played a significant role in understanding the development of the embryonic vasculature. Interestingly, zebrafish can survive the first week of development without a functional vasculature or heartbeat, unlike mammalian embryos which cannot survive without a functional cardiovascular system [15–18]. This viability allows us to perform detailed studies even in zebrafish with severe cardiovascular defects. In the zebrafish genome, approximately, 20% of the genes are duplicated (ref). For TF, zebrafish has two TF paralogs, designated as *f3a* and *f3b*[19], which are homologous to human and mouse TF genes [20] respectively. Knockdown of *f3b* using a morpholino oligonucleotide (MO) suggests that the *f3b* gene is important for coagulation and angiogenesis [21]. However, there are no reports on the role of *f3a* till date. In this study, we were able to successfully knockdown the *f3a* gene by MO and investigate the effects on bleeding and angiogenesis.

## Methods

### Phylogenetic analysis

TF protein and DNA sequences from human (*Homo sapiens*), rabbit (*Oryctolagus cuniculus*), mouse (*Mus musculus*), and zebrafish *(Danio rer*io) were retrieved from the National Center for Biotechnology Information (NCBI), European Molecular Biology Laboratory (EMBL), and Ensembl genome databases. Sequence alignment and construction of phylogenetic tree based on neighbor-joining method was performed using ClustalW2[22].

### Zebrafish strains and maintenance

The following zebrafish lines were used in this study: Tübingen (ZIRC [Zebrafish International Resource Center], Eugene, Oregon), *Tg(fli1a:nEGFP)y7 (Tg(gata1:dsRed)sd2*,[23] *and Tg(kdrl:mCherry-CAAX)y171[24]*. Adult fish were maintained under a constant temperature of 28.0°C and were subjected to 14-h light: 10-h ark photoperiod. All animal studies performed in these facilities were under the Medical College of Wisconsin–approved protocol for Animal Use Application. Freshly fertilized embryos were procured through natural breeding of adult zebrafish and were kept at 28.0°C in 1X E3 embryo medium (E3 medium) containing 5 mmol/L NaCl, 0.17 mmol/L KCl, 0.33 mmol/L CaCl_2_, 0.33 mmol/L MgSO_4_, and 0.05% methylene blue. In some instances, embryos were treated with 0.003% of 1-phenyl-2-thiourea (Sigma-Aldrich), starting at 24 hours postfertilization (hpf), to minimize pigmentation.

### Embryo collection and RNA extraction

Wild-type strain zebrafish were maintained at 23.5°C on a 14 h light/10 h dark cycle. At the time of mating, breeding males and females were separated overnight before letting them spawn naturally in the morning to allow for synchronization of developmental stages. Fertilized eggs were grown at 28.0°C and staged using previously defined criteria[25, 26]. Samples from 9 different developmental stages, 2 hpf to 3 days post-fertilization, were collected by snap freezing the embryos in dry ice.

RNA extraction was performed as described by Jong et al[27]. In brief, tubes with embryos from different timepoints were kept in liquid nitrogen and ground individually with a liquid nitrogen pre-chilled metal micro-pestle (Carl Roth). The pestle was lifted slightly and 200 μL Qiazol (Qiagen, USA) was added. The pestle was placed back into the tube with Qiazol and the homogenate was allowed to thaw. Before removal, the pestle was washed with an additional 100 μL Qiazol to rinse off any material that might have stuck to the pestle. The homogenate was vortexed vigorously for 30 seconds, left at room temperature for at least 5 minutes, and then spun down quickly for 30 seconds. 60 μL chloroform was added to the homogenate, vortexed for 15 seconds and kept at room temperature for 3 minutes. The partly separated mixture was transferred to a pre-prepared phase-lock gel heavy containing tube and centrifuged for 15 minutes at 12,000 × g. The aqueous phase was transferred to a new 1.5 mL tube. RNA was purified by column precipitation according to the RNeasy MinElute Cleanup kit (Qiagen, USA). At the end of the procedure, RNA was eluted in 12 μL nuclease-free water.

### *f3a* expression by RT-PCR

Double-stranded cDNA was synthesized using the QuantiTect Reverse transcription kit (Qiagen, USA). For RT-PCR analysis of *f3a*, the forward and reverse primers used were 5’- ACGTGGAGTCCAAAACCAAC -3’ and 5’- CAGCGCTGTAATAGGCCTTC -3’ and for Beta-actin (control), 5’- CGAGCTGTCTTCCCATCCA -3’ and 5’- TCACCAACGTAGCTGTCTTTCTG -3’. Transcript levels were quantified by SYBR™ Green PCR Master Mix (IDT, USA). For amplification by PCR, the initial denaturing step at 94°C for 5 minutes was followed by 40 amplification cycles of 30 seconds at 94°C; 30 seconds at 60°C; 60 seconds at 72°C, and a final extension period of 10 minutes at 72°C. *f3a* expression at different embryonic stages was calculated using beta-actin as a normalizing control and the ΔΔCt method.

### Morpholino approach

An antisense MO targeting the exon-intron junction (exon 3) of *f3a* (MO:*f3a*) was designed to knockdown the expression of *f3a*. As a negative control, we used a standard control MO (control-MO) specific to a human β-globin intron mutation. All MO solutions were synthesized by GeneTools (Oregon, USA). The MO sequences are as follows:

*f3a*-MO: 5’-AAACAACTAAACACTGACTGTCATG-3’
Control-MO: 5’-CCTCTTACCTCAGTTACAATTTATA-3’

All MO solutions were briefly heated at 65°C and resuspended in 1X Danieau buffer (58 mmol/L NaCl, 0.7 mmol/L KCl, 0.4 mmol/L MgSO4, 0.6 mmol/L Ca(NO_3_)_2_, 5.0 mmol/L HEPES, pH 7.6), and 0.1% (w/v) phenol red dye (Sigma-Aldrich, USA), to a final concentration of 8 ng/nL. Embryos at 1 to 2-cell stage were positioned in individual grooves made on a 1.0% agarose gel and were initially injected at concentrations ranging from 0.5 to 2 ng/nL.

### Screening of *f3a* knockdown by PCR and quantification

Total RNA extraction[27] and cDNA synthesis were performed from control- and *f3a*-MO injected embryos as described before[28]. For PCR-based gene expression analysis of *f3a*, the forward and reverse primers used were 5’-ACGTGGAGTCCAAAACCAAC-3’ and 5’-TCCGTCACATGCAGCTTTGT -3’; for *f3b*, 5’- CTTGGGGACCCAAACCTGTC -3’ and 5’- TCCAGTCGGTTAAACTCCGC -3’, and for GAPDH (control), 5’- GTGGAGTCTACTGGTGTCTTC -3’ and 5’- GTGCAGGAGGCATTGCTTACA -3’. Each amplification reaction was separated on a 1.5% agarose gel stained with ethidium bromide-stain and the bands were visualized using UV Transilluminator (Chemidoc System, USA). Gels were quantified using image J software[29].

### Hemoglobin staining

As described by Paffett-Lugassy and Zon[30], O-dianisidine staining was used to detect the presence of hemoglobin within intact 52 hpf zebrafish embryos injected with Control or *f3a* MOs. Dechorionated or hatched embryos were stained in the dark for 30 minutes at room temperature within a solution containing O-dianisidine (0.6 mg/ml), 0.01 M sodium acetate (pH 4.5), 0.65% H_2_O_2_, and 40% (vol/vol) ethanol. After staining, embryos were washed with RO water and then fixed in 4% paraformaldehyde (PFA) for 1 h at room temperature. Pigments were removed from fixed embryos by incubating in a solution of 0.8% KOH, 0.9% H_2_O_2_, and 0.1% Tween-20 for 30 minutes at room temperature. Embryos were then washed with phosphate-buffered saline (PBS) containing 0.1% Tween-20, and then fixed again in 4% PFA for 3 h before storage in PBS at 4°C. All embryos were positioned in dorsal recumbency and imaged using a Keyence BZ-X700 fluorescent microscope (Japan).

### Vascular development

Endothelial-specific transgenic lines of zebrafish (*flk1:egfp-NLS/kdrl:mCherry-CAAX*) were used to study vascular development. In brief, *Tg(fli1a:nEGFP)y7 (Tg(gata1:dsRed)sd2*,[23] *and Tg(kdrl:mCherry-CAAX)y171[24]* lines were maintained at 23.5°C on a 14 h light/10 h dark cycle. At the time of mating, breeding males and females were separated overnight before letting them spawn naturally in the morning. Embryos at 1 to 2-cell stage were positioned in individual grooves made on a 1.0% agarose gel and were initially injected at concentrations ranging from 1 to 2 ng/nL.

### Microscopy

For bright-field/fluorescent microscopy, wild-type or transgenic embryos were first embedded in 1.0% low melting agarose in E3 medium and 30 μg/mL tricaine mesylate. Embryos were mounted on 35 mm glass-bottom Petri dishes and imaged using Keyence BZ-X700 fluorescent microscope (Japan). A Texas Red filter cube (OP-87765, Keyence) was used to detect mCherry and dsRed (red fluorescent protein)-labeled cells, a GFP filter cube (OP-87763, Keyence) was used to detect GFP/EGFP-labeled tissues, and a 4’,6-diamidino-2-phenylindole (DAPI) filter cube (OP-87762, Keyence) was used to image DAPI-stained samples. Z-series bright-field and fluorescent images were acquired, and composite images were generated using the BZ-X Image Analyzer software. Brightness and contrast were adjusted using Fiji Software.

### Statistical analysis

Statistical analysis was performed using GraphPad Prism 8 software (GraphPad Software, USA) by Student’s t test (2 groups; nonparametric). Bleeding and morphological phenotypes were expressed as percentage differences from respective controls.

## Results

### Phylogenic analysis and protein identity of zebrafish TF-*f3b*

To understand the evolutionary relationships across different species, we compared the mouse, human and rabbit TF gene and protein sequences to the TF paralogs (*f3a* and *f3b*). Multiple sequence alignment (ClustalW) revealed that zebrafish *f3a* protein was more closely related to that of mouse and human compared with *f3b* (**Figure 1A**) and was more identical to human F3 compared to *f3b* (**Figure 1B**). RT-PCR analysis of expression during embryonic developmental stages showed that *f3a* was detected as early as 2h (64-cell stage) and in larvae at up to 36 hpf (**Figure 1C**). Previous report indicate detecting *f3a* transcripts at 64-cell stage, which suggest maternal contribution to larvae at this stage [26]. Between 2.75 hpf to 6 hpf, both maternal and zygotic expression appears to be present because *f3a* was strongly transcribed compared to earlier (2 hpf) and later stages (10, 18, and 24 hpf) **(Figure 1C)**. These observations indicate *f3a* transcription crossover from maternal to zygotic stages (between 2 hpf to 6 hpf). From 28 hpf to 36 hpf the expression of *f3a* was relatively stable, suggesting that *f3a* expression contributes to the development of yolk sac, circulation, pigmentation, heartbeat, circulation in the aortic arch, and fins [26]. At 48 hpf and 72 hpf expression was low compared to 36 hpf, suggesting *f3a* might not play a significant role in hatching and protruding mouth stages.

**Figure 1:**
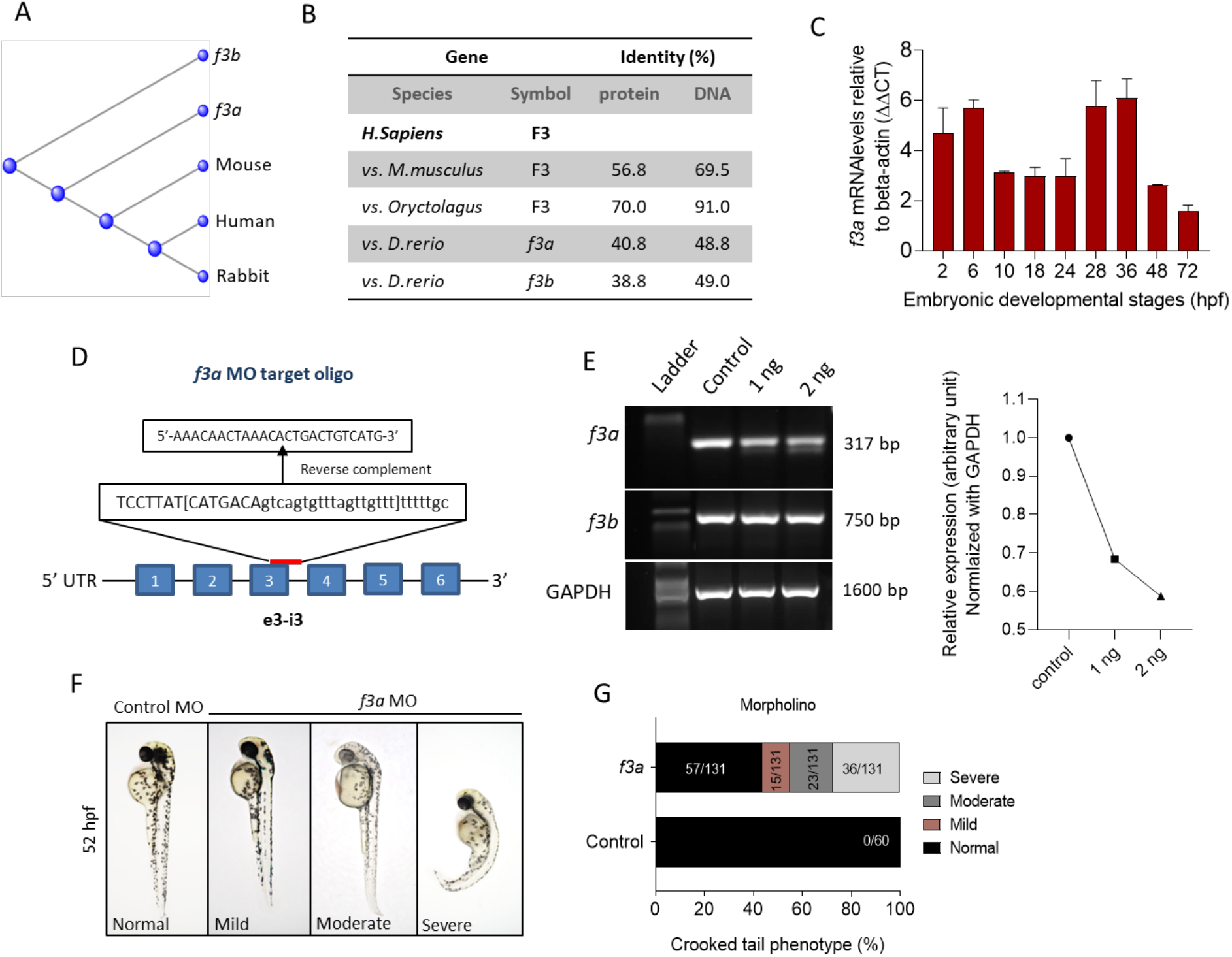
Evolutionary relationships and Spatiotemporal expression of Zebrafish TF-*f3b:* **(A)** Phylogenetic tree of tissue factor (F3) from zebrafish (Danio), mouse (Mus), rabbit (Oryc), and human (Homo). **(B)** Protein and DNA percent identity of F3 from zebrafish (Danio), mouse (Mus), rabbit (Oryc), and human (Homo). **(C)** mRNA expression from whole embryos demonstrates that zebrafish *f3a* is expressed in early embryonic stages. Beta-actin was used to normalize the expression of *f3a*. **(D)** A schematic diagram of *f3a* antisense morpholino oligonucleotide (MO) targeting the exon-intron junction (exon 3); design and synthesis by GeneTools (Oregon, USA). **(E)** The molecular targeting and efficiency of *f3a* MO-injected with 1 and 2 ng of *f3a* was assessed by PCR and quantified using ImageJ. **(F)** MO-mediated knockdown of *f3a* expression results in morphological abnormalities. Zebrafish embryos at 1-or 2-cell stage were injected with control and *f3a* MO and examined morphologically at 52 hpf. MO phenotypes: control MO injected (control MO), and mild, moderate, and severe phenotypes. **(G)** The morphological abnormalities were quantified. Numbers in the bars represent the ratios used to calculate the percentages. Data were pooled from three independent experiments.

### Spatiotemporal expression of zebrafish TF-*f3a*

Previously, Zhou and colleagues reported that *f3b* knockdown in zebrafish exhibits defective vasculogenesis [21]. Because f3a’s role during embryonic development was not known, we therefore examined its role by mRNA knockdown using MO antisense oligonucleotides (Gene Tools, USA) (**Figure 1D**). Embryos were collected freshly and 1 and 2 ng of *f3a* MO, targeted exon-intron junction (*e3-i3*), was injected at 1-to 2-cell stage. Twenty-four hours later, genomic DNA was isolated from the injected embryos and expression of *f3a* and *f3b* analyzed by PCR. Control group embryos were injected with a random sequence of standard MO. Our data revealed lower PCR transcript levels of *f3a*, but not *f3b*, at both 1 and 2 ng levels in injected embryos, suggesting specificity of the targeted knockdown towards *f3a* **(Figure 1E)**.

### *f3a* knockdown by Morpholinos approach and morphological characterization

The *f3a* morphants, injected with 2 ng of MO, were examined at 52 hpf for any defect in morphological appearance. Control MO-injected embryos appeared to be normal, with properly developed head and eyes, and elongated body axis, and a long tail **(Figure 1F)**. In contrast, embryos injected with 2 ng of *f3a* MO exhibited a shortened body axis and a short, crooked tail. Based on the degree of severity, we segregated them as mild, moderate, and severe and quantified the severity. On an average, 11.45% (15/131) showed mild phenotype with shortened body axis. Moderate defect in morphological appearance was observed in 17.55% (23/131), with a partially curved tail. Severe morphological defect was observed in 27.5% (36/131) of embryos, with curved body axis and defectively developed tail **(Figure 1G)**.

### Bleeding phenotype in *f3a* knockdown zebrafish

As TF is a triggering factor for activation and coagulation cascade, in a separate experiment, we checked whether the knockdown of *f3a* showed bleeding phenotype. We observed embryos at 52 hpf, which showed bleeding mostly in the head and tail regions **(Figure 2A)**. Control MO-injected embryos appeared normal with no visible bleeding. Conversely, *f3a* MO-injected embryos showed 5.44% (8/147) and 9.52% (14/147) bleeding in tail and head regions, respectively **(Figure 2B)**. With the next set of *f3a* and standard MO injection, at 52 hpf, we stained the embryos with O-dianisidine to detect hemoglobin-containing cells **(Figure 2C)**; around 7.5% had bleeding in the head. No bleeding was observed in control MO-injected embryos at 52 hpf **(Figure 2C,D)**. Common cardinal vein (CCV) region also stained with O-dianisidine due to the presence of a pool of blood cells and heart. This region appears stained irrespective of phenotype, and therefore, we did not consider the CCV region for bleeding score. These findings confirm that, similar to *f3b [21], f3a* plays a role in vascular stability in our zebrafish model.

**Figure 2:**
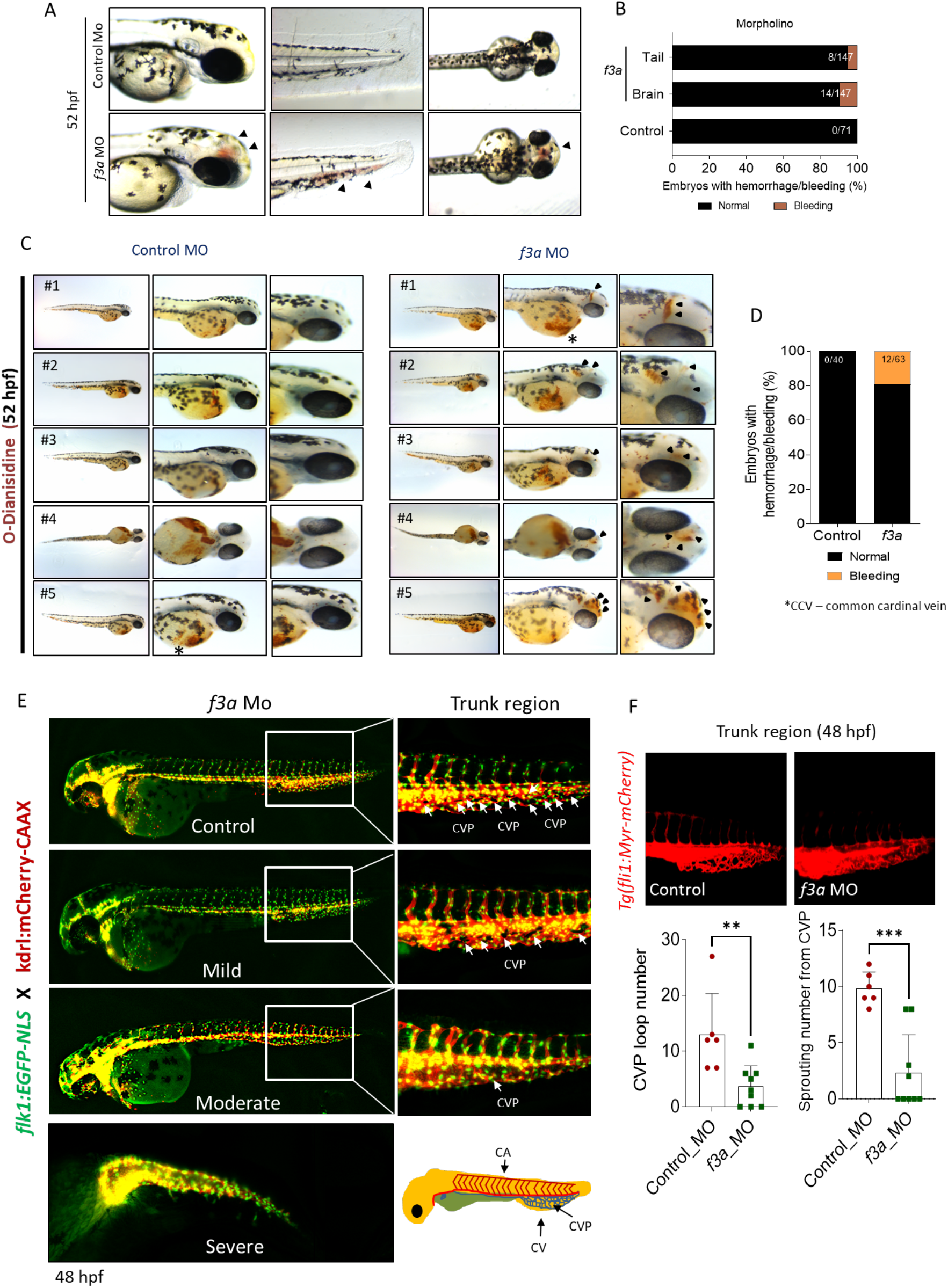
*f3a* knockdown showed bleeding and delayed angiogenesis. **(A)** Lateral and dorsal views of head, and lateral view of tail in live embryos. The arrows depict brain and tail bleeding at 52 hpf and **(B)** quantification of the same. Numbers in the bars represent the ratios used to calculate the percentages. **(C)** Bright-field images of zebrafish embryos stained with O-dianisidine (OD) and imaged at 48 hpf. Black arrows indicate sites where abnormal accumulation of hemoglobinized blood was detected in the head region (control MO and *f3a* MO). **(D)** Percentage of cerebral hemorrhage in embryos at 52 hpf. Numbers in the bars represent the ratios used to calculate the percentages. Data were pooled from three independent experiments. **(E)** To study vascular development, endothelial-specific transgenic lines of zebrafish (*flk1:egfp-NLS/kdrl:mCherry-CAAX*) embryo were used to knockdown *f3a*. Images represent MO-mediated knockdown of *f3a* expression resulting in morphological abnormalities. MO phenotypes: control MO injected (control MO), and mild, moderate, and severe phenotypes. Differences in caudal vein plexus (CVP) in the trunk regions are indicated by white arrows. **(F)** *Tg(fli1:Myr-mCherry)* line was used to quantify CVP loop number and Sprouting number from the CVP at 48 hpf. Embryos were randomly picked from ~60 MO-injected (6 from control and 9 from *f3a* MO) embryos from control and *f3a*.

### Defective angiogenesis in *f3a* knockdown zebrafish

Next, we investigated the effect of *f3a* knockdown on vasculogenesis and angiogenesis using endothelial-specific transgenic lines of zebrafish (*flk1:egfp-NLS/kdrl:mCherry-CAAX*) and imaged vascular development. Interestingly, we did not observe any defect in vessel formation at 48 hpf between control and *f3a*-MO-injected groups **(Figure 2E)**, suggesting that *f3a* might not be involved in initial development of central aorta and no major deficit in vasculogenesis. During zebrafish embryonic angiogenesis, caudal vein plexus (CVP) and inter-segmental vessel (ISV) formation are the two obvious vessel patterns observed. In *f3a* knockdown embryos, the development of CVP was greatly abrogated **(Figure 2E)**. Quantification of loop formation at mean CVP showed a 3-fold decrease in *f3a* morphants versus controls at 48 hpf **(Figure 2F)**. The CVP forms during a very active period of angiogenic sprouting. Thus, we looked at the difference in the CVP angiogenic sprouting at 48 hpf. Like number of CVP, the mean spouting from CVP was 4-fold lower in *f3a* injected MO than the control MO-injected group **(Figure 2F)**. Occasionally, we also observed a defect in central arteries (CtAs) and primordial midbrain channel (PMBC) in *f3a* morphant CVP defective embryos. However, we did not see any impairment in the intersegmental vessel (ISV) development. These findings support the notion that *f3a is* required for angiogenesis.

## Discussion

In the present study, we used a zebrafish knockdown model to study one of the homologs genes of TF (*f3a*) to gain insights into the developmental aspects. Previously, Zhou and his colleagues investigated the role of *f3b* in zebrafish by morpholino approach and reported that *f3b* knockdown in zebrafish exhibits defective vasculogenesis[21]. In this study, we successfully knocked down *f3a* by morpholino approach and investigated its effects on bleeding and vascular development.

Phylogenetic analysis revealed that *f3a* is more closely related to humans and mice than *f3b*. Spatiotemporal expression of *f3a* by RT-PCR revealed that unlike *f3b* (expressed only after 18 hpf), *f3a* expression occurs across all the stages, suggesting that *f3a* transcripts are both maternally and zygotically expressed. Expression of *f3a* is stable throughout the embryonic stages. *f3a* MO-injected embryos showed morphological abnormalities, including shorter body lengths and bent tails. The standard MO-injected embryos had no phenotype changes.

Evaluating the vascular development using endothelial-specific transgenic lines of zebrafish (*flk1:egfp-NLS/kdrl:mCherry-CAAX*) revealed early vascular development (central aorta and ISV) remain stable even after *f3a* knockdown. Interestingly, the development of CVP, which is composed of a dorsal and ventral vein with interconnecting vessels, and spouting from CVP were suppressed in the zebrafish *f3a* knockdown models, signifying the contribution of *f3a* in angiogenesis. Despite defective angiogenesis, knockdown of *f3a* also resulted in mild bleeding in the brain and tail regions. Similar bleeding characteristics were observed in zebrafish *f3b* knockdown models[21].

In conclusion, similar to *f3b* knockdown[21], *f3a* knockdown also showed a mild bleeding phenotype, suggesting that both *f3a* and *f3b* are required for effective hemostasis function in zebrafish. Our study also confirmed that *f3a* is essential for angiogenesis in addition to its involvement in hemostasis. In addition, although we see a stable formation of central aorta, which implies no defect in vasculogenesis, at 48 hpf in control and *f3a*-MO-injected groups, it must be further validated at earlier time points (12-18 hpf). Nevertheless, a CRISPR/cas9-mediated gene knockout approach would provide more insights (molecular and signaling importance) to understand the non-hemostasis function of TF paralogs (*f3a* and *f3b*) in zebrafish.

## Acknowledgements

This work was supported by a grant from CSL Behring (Prof. Heimburger award to S.S.).

R. R. is supported by R61 HL154254-01, and from programmatic development funds from Department of Pediatrics, and Children’s Research Institute, Milwaukee, WI. The authors thank Dr. Shahram Eisa-Beygi for the initial technical support.

## Authorship

S. S. performed the experiments and analyzed data. R.R., H.W., J.L., and C.F., provided expert technical advice and contributed to the study and edited the manuscript. S.S. designed the experiment, analyzed data, and wrote the manuscript.

## References

[1] N. Mackman, J.H. Morrissey, B. Fowler, T.S. Edgington, Complete sequence of the human tissue factor gene, a highly regulated cellular receptor that initiates the coagulation protease cascade, Biochemistry 28(4) (1989) 1755–62.

[2] J.H. Morrissey, H. Fakhrai, T.S. Edgington, Molecular cloning of the cDNA for tissue factor, the cellular receptor for the initiation of the coagulation protease cascade, Cell 50(1) (1987) 129–35.

[3] E.K. Spicer, R. Horton, L. Bloem, R. Bach, K.R. Williams, A. Guha, J. Kraus, T.C. Lin, Y. Nemerson, W.H. Konigsberg, Isolation of cDNA clones coding for human tissue factor: primary structure of the protein and cDNA, Proc Natl Acad Sci U S A 84(15) (1987) 5148–52.

[4] R. Pawlinski, N. Mackman, Tissue factor, coagulation proteases, and protease-activated receptors in endotoxemia and sepsis, Critical care medicine 32(5 Suppl) (2004) S293–7.

[5] D. Chen, A. Dorling, Critical roles for thrombin in acute and chronic inflammation, Journal of thrombosis and haemostasis: JTH 7 Suppl 1 (2009) 122–6.

[6] J. Erlich, G.C. Parry, C. Fearns, M. Muller, P. Carmeliet, T. Luther, N. Mackman, Tissue factor is required for uterine hemostasis and maintenance of the placental labyrinth during gestation, Proc Natl Acad Sci U S A 96(14) (1999) 8138–43.

[7] P. Carmeliet, N. Mackman, L. Moons, T. Luther, P. Gressens, I. Van Vlaenderen, H. Demunck, M. Kasper, G. Breier, P. Evrard, M. Muller, W. Risau, T. Edgington, D. Collen, Role of tissue factor in embryonic blood vessel development, Nature 383(6595) (1996) 73–5.

[8] J.R. Toomey, K.E. Kratzer, N.M. Lasky, J.J. Stanton, G.J. Broze, Jr., Targeted disruption of the murine tissue factor gene results in embryonic lethality, Blood 88(5) (1996) 1583–7.

[9] T.H. Bugge, Q. Xiao, K.W. Kombrinck, M.J. Flick, K. Holmback, M.J. Danton, M.C. Colbert, D.P. Witte, K. Fujikawa, E.W. Davie, J.L. Degen, Fatal embryonic bleeding events in mice lacking tissue factor, the cell-associated initiator of blood coagulation, Proc Natl Acad Sci U S A 93(13) (1996) 6258–63.

[10] E.G. Tuddenham, S. Pemberton, D.N. Cooper, Inherited factor VII deficiency: genetics and molecular pathology, Thrombosis and haemostasis 74(1) (1995) 313–21.

[11] J. Zhang, J. Ding, X. Zhang, X. Shao, Z. Hao, Regulation of vascular endothelial growth factor (VEGF) production and angiogenesis by tissue Factor (TF) in SGC-7901 gastric cancer cells, Cancer Biol Ther 4(7) (2005) 769–72.

[12] M. Belting, M.I. Dorrell, S. Sandgren, E. Aguilar, J. Ahamed, A. Dorfleutner, P. Carmeliet, B.M. Mueller, M. Friedlander, W. Ruf, Regulation of angiogenesis by tissue factor cytoplasmic domain signaling, Nat Med 10(5) (2004) 502–9.

[13] J. Chen, A. Bierhaus, S. Schiekofer, M. Andrassy, B. Chen, D.M. Stern, P.P. Nawroth, Tissue factor--a receptor involved in the control of cellular properties, including angiogenesis, Thrombosis and haemostasis 86(1) (2001) 334–45.

[14] F. Tonelli, J.W. Bek, R. Besio, A. De Clercq, L. Leoni, P. Salmon, P.J. Coucke, A. Willaert, A. Forlino, Zebrafish: A Resourceful Vertebrate Model to Investigate Skeletal Disorders, Front Endocrinol (Lausanne) 11 (2020) 489.

[15] D.Y. Stainier, B.M. Weinstein, H.W. Detrich, 3rd, L.I. Zon, M.C. Fishman, Cloche, an early acting zebrafish gene, is required by both the endothelial and hematopoietic lineages, Development 121(10) (1995) 3141–50.

[16] S.W. Jin, W. Herzog, M.M. Santoro, T.S. Mitchell, J. Frantsve, B. Jungblut, D. Beis, I. C. Scott, L.A. D’Amico, E.A. Ober, H. Verkade, H.A. Field, N.C. Chi, A.M. Wehman, H. Baier, D.Y. Stainier, A transgene-assisted genetic screen identifies essential regulators of vascular development in vertebrate embryos, Dev Biol 307(1) (2007) 29–42.

[17] D.Y. Stainier, Zebrafish genetics and vertebrate heart formation, Nat Rev Genet 2(1) (2001) 39–48.

[18] A.J. Sehnert, A. Huq, B.M. Weinstein, C. Walker, M. Fishman, D.Y. Stainier, Cardiac troponin T is essential in sarcomere assembly and cardiac contractility, Nat Genet 31(1) (2002) 106–10.

[19] K. Howe, M.D. Clark, C.F. Torroja, J. Torrance, C. Berthelot, M. Muffato, J.E. Collins, S. Humphray, K. McLaren, L. Matthews, S. McLaren, I. Sealy, M. Caccamo, C. Churcher, C. Scott, J.C. Barrett, R. Koch, G.J. Rauch, S. White, W. Chow, B. Kilian, L.T. Quintais, J. A. Guerra-Assuncao, Y. Zhou, Y. Gu, J. Yen, J.H. Vogel, T. Eyre, S. Redmond, R. Banerjee, J. Chi, B. Fu, E. Langley, S.F. Maguire, G.K. Laird, D. Lloyd, E. Kenyon, S. Donaldson, H. Sehra, J. Almeida-King, J. Loveland, S. Trevanion, M. Jones, M. Quail, D. Willey, A. Hunt, J. Burton, S. Sims, K. McLay, B. Plumb, J. Davis, C. Clee, K. Oliver, R. Clark, C. Riddle, D. Elliot, G. Threadgold, G. Harden, D. Ware, S. Begum, B. Mortimore, G. Kerry, P. Heath, B. Phillimore, A. Tracey, N. Corby, M. Dunn, C. Johnson, J. Wood, S. Clark, S. Pelan, G. Griffiths, M. Smith, R. Glithero, P. Howden, N. Barker, C. Lloyd, C. Stevens, J. Harley, K. Holt, G. Panagiotidis, J. Lovell, H. Beasley, C. Henderson, D. Gordon, K. Auger, D. Wright, J. Collins, C. Raisen, L. Dyer, K. Leung, L. Robertson, K. Ambridge, D. Leongamornlert, S. McGuire, R. Gilderthorp, C. Griffiths, D. Manthravadi, S. Nichol, G. Barker, S. Whitehead, M. Kay, J. Brown, C. Murnane, E. Gray, M. Humphries, N. Sycamore, D. Barker, D. Saunders, J. Wallis, A. Babbage, S. Hammond, M. Mashreghi-Mohammadi, L. Barr, S. Martin, P. Wray, A. Ellington, N. Matthews, M. Ellwood, R. Woodmansey, G. Clark, J. Cooper, A. Tromans, D. Grafham, C. Skuce, R. Pandian, R. Andrews, E. Harrison, A. Kimberley, J. Garnett, N. Fosker, R. Hall, P. Garner, D. Kelly, C. Bird, S. Palmer, I. Gehring, A. Berger, C.M. Dooley, Z. Ersan-Urun, C. Eser, H. Geiger, M. Geisler, L. Karotki, A. Kirn, J. Konantz, M. Konantz, M. Oberlander, S. Rudolph-Geiger, M. Teucke, C. Lanz, G. Raddatz, K. Osoegawa, B. Zhu, A. Rapp, S. Widaa, C. Langford, F. Yang, S.C. Schuster, N.P. Carter, J. Harrow, Z. Ning, J. Herrero, S.M. Searle, A. Enright, R. Geisler, R.H. Plasterk, C. Lee, M. Westerfield, P.J. de Jong, L.I. Zon, J.H. Postlethwait, C. Nusslein-Volhard, T.J. Hubbard, H. Roest Crollius, J. Rogers, D.L. Stemple, The zebrafish reference genome sequence and its relationship to the human genome, Nature 496(7446) (2013) 498–503.

[20] C. Stein, M. Caccamo, G. Laird, M. Leptin, Conservation and divergence of gene families encoding components of innate immune response systems in zebrafish, Genome biology 8(11) (2007) R251.

[21] R.F. Zhou, Y. Liu, Y.X. Wang, W. Mo, M. Yu, Coagulation factor III (tissue factor) is required for vascularization in zebrafish embryos, Genetics and molecular research: GMR 10(4) (2011) 4147–57.

[22] J.H. Hung, Z. Weng, Sequence Alignment and Homology Search with BLAST and ClustalW, Cold Spring Harb Protoc 2016(11) (2016).

[23] S. Eisa-Beygi, F.M. Benslimane, S. El-Rass, S. Prabhudesai, M.K.A. Abdelrasoul, P.M. Simpson, H.C. Yalcin, P.E. Burrows, R. Ramchandran, Characterization of Endothelial Cilia Distribution During Cerebral-Vascular Development in Zebrafish ( Danio rerio), Arterioscler Thromb Vasc Biol 38(12) (2018) 2806–2818.

[24] A. Borovina, S. Superina, D. Voskas, B. Ciruna, Vangl2 directs the posterior tilting and asymmetric localization of motile primary cilia, Nat Cell Biol 12(4) (2010) 407–12.

[25] A.S. Machikhin, M.V. Volkov, A.B. Burlakov, D.D. Khokhlov, A.V. Potemkin, Blood Vessel Imaging at Pre-Larval Stages of Zebrafish Embryonic Development, Diagnostics (Basel) 10(11) (2020).

[26] C.B. Kimmel, W.W. Ballard, S.R. Kimmel, B. Ullmann, T.F. Schilling, Stages of embryonic development of the zebrafish, Dev Dyn 203(3) (1995) 253–310.

[27] M. de Jong, H. Rauwerda, O. Bruning, J. Verkooijen, H.P. Spaink, T.M. Breit, RNA isolation method for single embryo transcriptome analysis in zebrafish, BMC Res Notes 3 (2010) 73.

[28] S.M. Peterson, J.L. Freeman, RNA isolation from embryonic zebrafish and cDNA synthesis for gene expression analysis, J Vis Exp (30) (2009).

[29] C.A. Schneider, W.S. Rasband, K.W. Eliceiri, NIH Image to ImageJ: 25 years of image analysis, Nat Methods 9(7) (2012) 671–5.

[30] N.N. Paffett-Lugassy, L.I. Zon, Analysis of hematopoietic development in the zebrafish, Methods Mol Med 105 (2005) 171–98.

